# Biology and Bias in Cell Type-Specific RNAseq of Nucleus Accumbens Medium Spiny Neurons

**DOI:** 10.1101/444315

**Authors:** Hope Kronman, Felix Richter, Benoit Labonté, Ramesh Chandra, Shan Zhao, Gabriel Hoffman, Mary Kay Lobo, Eric E. Schadt, Eric J. Nestler

## Abstract

Isolation of cell populations is untangling complex biological interactions, but studies comparing methodologies lack in vivo complexity and draw limited conclusions about the types of transcripts identified by each technique. Furthermore, few studies compare FACS-based techniques to ribosomal affinity purification, and none do so genome-wide. We addressed this gap by systematically comparing nuclear-FACS, whole cell-FACS, and RiboTag affinity purification in the context of D1 or D2 dopamine receptor-expressing medium spiny neuron (MSN) subtypes of the nucleus accumbens (NAc), a key brain reward region. We find that nuclear-FACS-seq generates a substantially longer list of differentially expressed genes between these cell types, and a significantly larger number of neuropsychiatric GWAS hits than the other two methods. RiboTag-seq has much lower coverage of the transcriptome than the other methods, but very efficiently distinguishes D1- and D2-MSNs. We also demonstrate differences between D1- and D2-MSNs with respect to RNA localization, suggesting fundamental cell type differences in mechanisms of transcriptional regulation and subcellular transport of RNAs. Together, these findings guide the field in selecting the RNAseq method that best suits the scientific questions under investigation.

## Introduction

The ability to isolate individual populations of cells with a homogeneous molecular profile has allowed biological inquiry to address new, more refined regulatory functions. Studies are parsing biological signals from complex, heterogeneous tissues and thereby revealing effects masked by cell type variability^1–4^. This work brings us one step closer to understanding the quantum nature of biology and to developing therapeutics that leverage its heterogeneity, rather than ones that lose efficacy as a consequence of it. We report here a comprehensive, technical comparison of cell isolation methodologies for RNAseq, and leverage the biological context of our study to provide new insight into the function of D1- and D2-type medium spiny neurons (MSNs) of the nucleus accumbens (NAc), part of the ventral striatum.

While techniques to isolate cell populations have been widely used in various tissues, they have not been systematically compared in a well-controlled framework for brain. There are substantial differences between methods; for example, whole cell-FACS and RiboTag extract cells from fresh tissue, while nuclear-FACS does so typically from frozen tissue; RiboTag and nuclear-FACS use a mechanical dissociation protocol, while whole cell-FACS uses an enzymatic one; and each recovers a different subpopulation of RNA species. Based on the literature and on procedural documentation for these methods, we hypothesize that they uncover related but only partially overlapping gene sets and biological functions.

Unlike library preparation methods for whole-genome and whole-transcriptome analysis, which have been carefully compared and assessed for biasing effects^5–8^, cell isolation protocols have been inconsistently compared and have yielded conflicting results. Several studies have analyzed differences generated by isolation of either whole cells or nuclei, but these studies are often performed in cell culture^9,10^, which lacks the heterogeneity and variability of in vivo tissues, and they draw limited conclusions about the biological types of transcripts isolated by each technique. Comparative studies on more complex tissues confine their interpretation to effects of each technique on a subset of differentially expressed genes, transcript length or RNA biotype^11^.

Few RNAseq studies have included a comparison between FACS-isolated cells/nuclei and ribosomal purification techniques like TRAP or RiboTag, and therefore ignore the important question about which RNA species are isolated by the latter methods that capture active translation^12–14^. Those studies that include such a comparison have relatively simple biological endpoints like method repeatability and contamination, and lack the comprehensiveness of a whole-genome analysis^10,15,16^.

The absence of a methodical comparison of whole cell-FACS, nuclear-FACS, and RiboTag affinity purification is becoming increasingly problematic as an increasing number of studies using these techniques are published and their results taken at face value. These techniques capture different cellular processes while simultaneously defining the same cellular identity. Only a head-to-head comparison for the same cell types can demonstrate if they predominantly capture differences or similarities, and identify the nature of those biases. The present study was designed to address this deficiency in the field.

Our two cell types of choice are both principal GABAergic MSNs of the NAc, a forebrain region implicated in reward and motivation. Both MSN subtypes respond to dopamine, but do so through the activity of different dopamine receptors^17,18^, display different physiology in response to reward-related stimuli^19–21^, and generate different behavioral outcomes^22–29^. Whole cell-FACS^17^ and TRAP^12^ have been used to distinguish between D1- and D2-MSNs, but the two methods have not been compared directly and these prior studies focused on the entire striatal complex of which the NAc represents a small sub-region. Here, we use all three RNA isolation methods – whole cell-FACS, nuclear-FACS, and RiboTag – to provide a deeper characterization of these behaviorally relevant NAc cell types than ever before. Most importantly, this study presents the first genome-wide, biological network-focused analysis of the contribution of these three methods to RNA characterization.

## Results

### Comparison of library complexity and distribution

As noted in the Introduction, whole cell-FACS, nuclear-FACS, and RiboTag affinity purification have methodological differences and retrieve different subcellularly-located RNAs (Fig. 1a). To compare the ability of these methods to distinguish D1- and D2-MSN populations from the NAc, we generated RNAseq libraries using ribo-depleted, total RNA isolated from the NAc of individual D1- or D2-Cre mice (see Methods). The libraries were prepared using the same kit and sequenced on the same platform using the same parameters. Thirty-nine samples were used for downstream analyses, constituting 16 whole cell-FACS (D1 n=9, D2 n=7), 11 nuclear-FACS (D1 n=6, D2 n=5), and 12 RiboTag (D1 n=6, D2 n=6) samples. We first confirmed that nuclear sequencing produces a larger percentage of intronic reads (Supplementary Fig. 1a), consistent with previous findings^30,31^.

**Figure 1.**
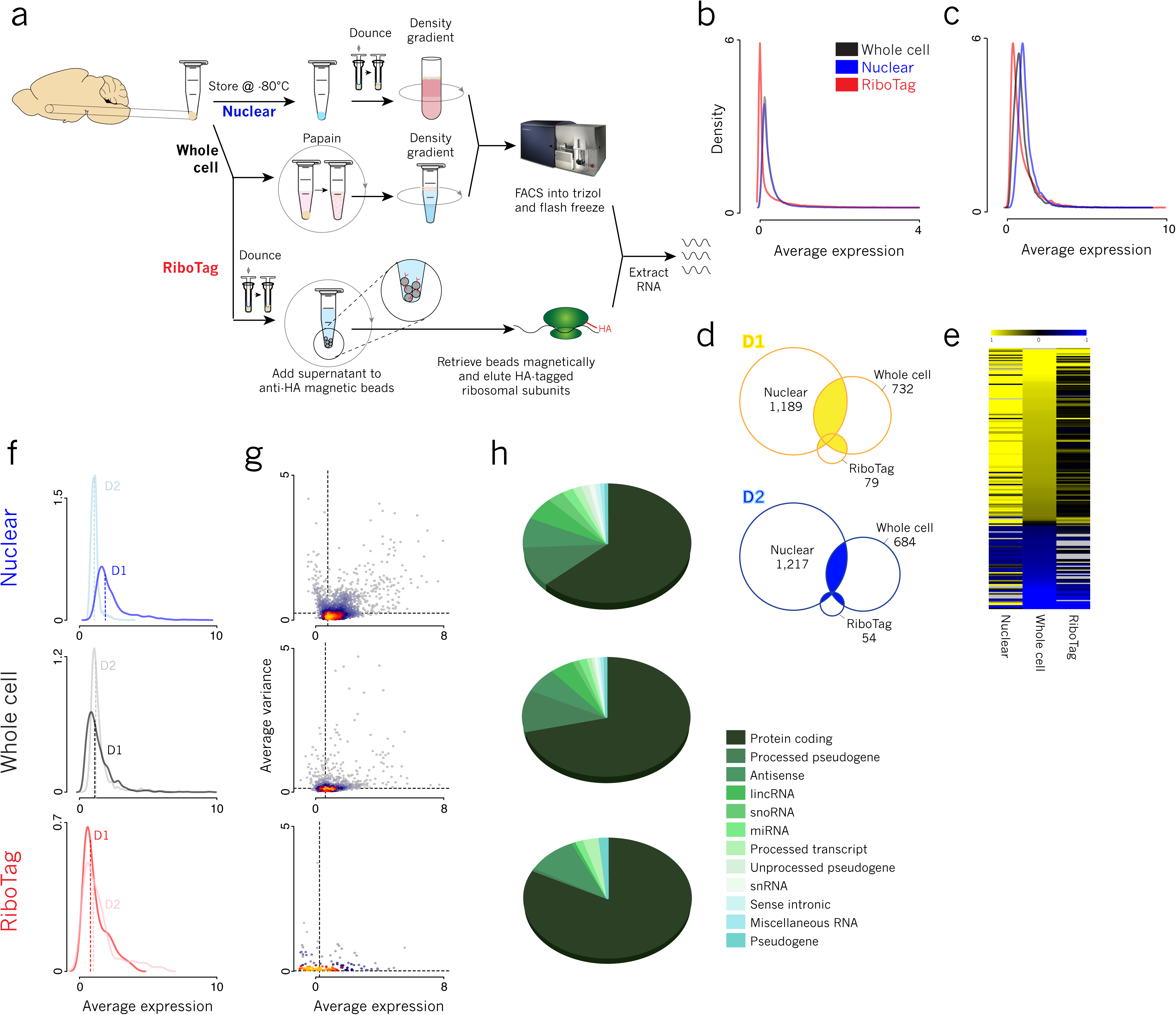
Library characterization demonstrates fewer differentially expressed transcripts and a predominance of protein coding genes in RiboTag compared to whole cell and nuclear RNAseq. (A) Method scheme for whole cell FACS, nuclear FACS, and RiboTag affinity purification showing key differences in steps involving sample preparation, cell dissociation, and retrieval of cellular fractions (B) Density plot for all methods of ln([average RPKM] +1) of all genes captured shows that RiboTag does not capture a number of genes (C) Density plots for all methods of ln([average RPKM] +1) of differentially expressed genes shows similar distributions across method (D) Overlap of D1- and D2-enriched differentially expressed genes across all methods (total D1 overlap = 134; total D2 overlap = 64) (E) Fold change of differentially expressed genes from most D1-enriched in yellow to most D2-enriched in blue, sorted by fold change in whole cell (black = low fold change, grey = not detected in the dataset) (F) Density plots for each method of ln([average RPKM] +1) of D1 (dark line) and D2 (light line) differentially expressed genes in the respective cell types; medians are indicated with dashed lines (G) Mean-variance plots comparing ln(variance) to ln([average RPKM] +1) of differentially expressed genes in pooled D1- and D2-MSNs show only slight differences across methods (H) Gene biotype distributions for each method’s differentially expressed genes show a decreasing proportion of protein coding genes from the RiboTag to the whole cell to the nuclear dataset

Differential RNA expression between D1 and D2 MSNs was performed within each method, and signatures were compared across methods. Using standard FPM (≥1 in ≥2 samples) and differential expression filtering (pAdj ≤ 0.05, 30% change, top 75% of mean size factor normalized counts), overlap in differentially expressed genes (DEGs) between conditions was not very high (Supplementary Fig, 1b), and the nuclear DEG pool was large and non-specific. Based on this nuclear DEG observation, we implemented a strict filter to remove genes with very low and very high value FPKMs (keeping genes with FPKM ≥ 1 in at least one sample and ≤ 5×10^4^ in all samples, Supplementary Fig. 1c). These filters reduced intronic and intergenic noise^32^ and eliminated artifacts of sample-specific overamplification of short-length genes. In contrast to nuclear and whole cell libraries, the RiboTag method had more lowly-expressed genes (Fig. 1b), necessitating different filtering cut-offs. This is consistent with previous literature^33^. We therefore implemented a less strict low-FPKM filter for the RiboTag dataset balanced by requiring observations in multiple samples (FPKM ≥0.1 in ≥2 samples). This filter dramatically increased RiboTag overlap compared to that found using the stringent FPKM filter for all methods (Supplementary Fig. 1d). These empirically-determined FPKM filtering parameters optimized comparison between the conditions, generating similar transcriptome-wide FPKM distributions across method (Fig. 1c).

Nuclear DEGs appeared to still be somewhat non-specific, and we ran a logistic regression to determine parameters contributing to nuclear DEG overlap vs. non-overlap (Supplementary Table 1). Perhaps due to increased variance in the set of nuclear DEGs (Fig. 1d), StandardError(log_2_foldchange) was significantly higher in non-overlapping nuclear DEGs compared to overlapping ones (logistic regression β = 5.4, p = 2.0 x 10^-2^). We therefore removed nuclear DEGs within the top quartile of StandardError(log_2_foldchange), greatly improving the specificity of this pool (Fig. 1d), and generating a robust population of cross-method D1-MSN- and D2-MSN-enriched genes, whose fold change followed similar patterns across method (Fig. 1e)

More genes were differentially expressed between the two cell types in the nuclear (2,361) compared to the whole cell (1,416) and RiboTag (133) datasets (Fig. 1d, Supplementary Table 2). These nuclear DEGs included genes of a wide variety of coding and non-coding biotypes (Fig. 1g), and the percentage of protein-coding DEGs increased steadily from nuclear (61.59%) to whole cell (69.52%) to RiboTag (82.71%). This bias towards protein-coding genes in the RiboTag dataset manifests before the implementation of differential expression, as demonstrated by the method’s relatively lower capture of short-length genes across the transcriptome, an effect which disappears when analyzing only protein-coding genes (Supplementary Fig. 1e)

Density plots of whole cell and RiboTag DEG average FPKMs within D1-MSNs and D2-MSNs did not show a detectable difference in distribution between cell types (Fig. 1e, Supplementary Table 3). However, in the nuclear dataset, there was a notable difference between cell types, with D2-enriched DEGs having a lower average FPKM than D1-enriched DEGs. Given its lower expression level and small overlap with RiboTag D2-DEGs, the D2 nuclear profile may represent stochastic transcriptional noise in D2-MSNs which, at baseline, does not correspond to active translation, but which poises cells to respond to future stimuli.

Nuclear DEGs had a higher median log(FPKM + 1) (0.76) than whole cell (0.61) or RiboTag (0.24) DEGs, as well as a higher median variance (nuclear = 0.24, whole cell = 0.13, RiboTag = 0.02) (Fig. 1f). This is largely driven by the higher median variance of nuclear D2-DEGs (0.31) than nuclear D1-DEGs (0.21). Importantly, within genes passing the FPKM filter in all conditions, FPKM is significantly correlated across method, as is variance (Supplementary Table 4).

Finally, we demonstrate that these data can be used to model RNA regulatory dynamics across cellular compartments, extending previous work with ribosomal and whole cell data^34^. Using a negative binomial generalized linear model (Supplementary Fig. 2a), effects of transcriptional, cytosolic and translational regulation were deduced. Genes undergoing one or more of these levels of regulation were overlapped (Supplementary Fig. 2b). The largest overlap was seen for genes transcriptionally enriched in D1 nuclei, but cytosolically enriched in D2-MSNs. Upstream miRNA analysis of this list using miRTarBase^35^ generated 27 mouse miRNAs (Fisher’s exact test two-sided adjusted p-value < 0.05), and 4 of these were differentially enriched in D2-nuclei but not in D2-whole cells (Supplementary Fig. 2c), pointing to cell type-specific mechanisms of subcellular transcriptional regulation. Numbers of genes falling into each category of regulation suggest that D1-MSNs undergo predominantly transcriptional and translational regulation, while D2-MSNs undergo predominantly cytosolic regulation.

### Variance partitioning of the combined datasets

Having assessed the transcriptome-coverage and basic library composition for each method, we analyzed the major sources of variance between and within the three datasets. A principal component analysis revealed strong separation of samples by method (Fig. 2a), with RiboTag samples differing the most from the other two methods, and clustering together most tightly. The Euclidean distances between Method cluster centroids were higher for RiboTag (*d*(WholeCell,Nuclear)=41, *d*(RiboTag,Nuclear)=75, *d*(RiboTag,WholeCell)=69) and the mean Euclidean distances within Method to each cluster centroid were significantly smaller for RiboTag (*d̄*_RiboTag_=1.7, *d̄*_WholeCell_=5.9, *d̄*_Nuclear_=10.9, two-sided t-test p<5×10^−5^ for both RiboTag comparisons, p=0.005 for WholeCell vs Nuclear). Samples from each method separated clearly by cell type (Fig. 2a, insets), and this was also seen in PC5/PC6 of the combined data (Supplementary Fig. 3a). Method accounted for more variability than cell type, though this is likely due to the similarity of the two cell types studied. When comparing our datasets to published RiboTag RNAseq from liver cells^33^, most of the variability in the combined data (PC1, 50.5%) was due to tissue, while less variability (PC2, 30.6%) separated RiboTag datasets from whole cell and nuclear (Supplemental Fig. 3b)

**Figure 2.**
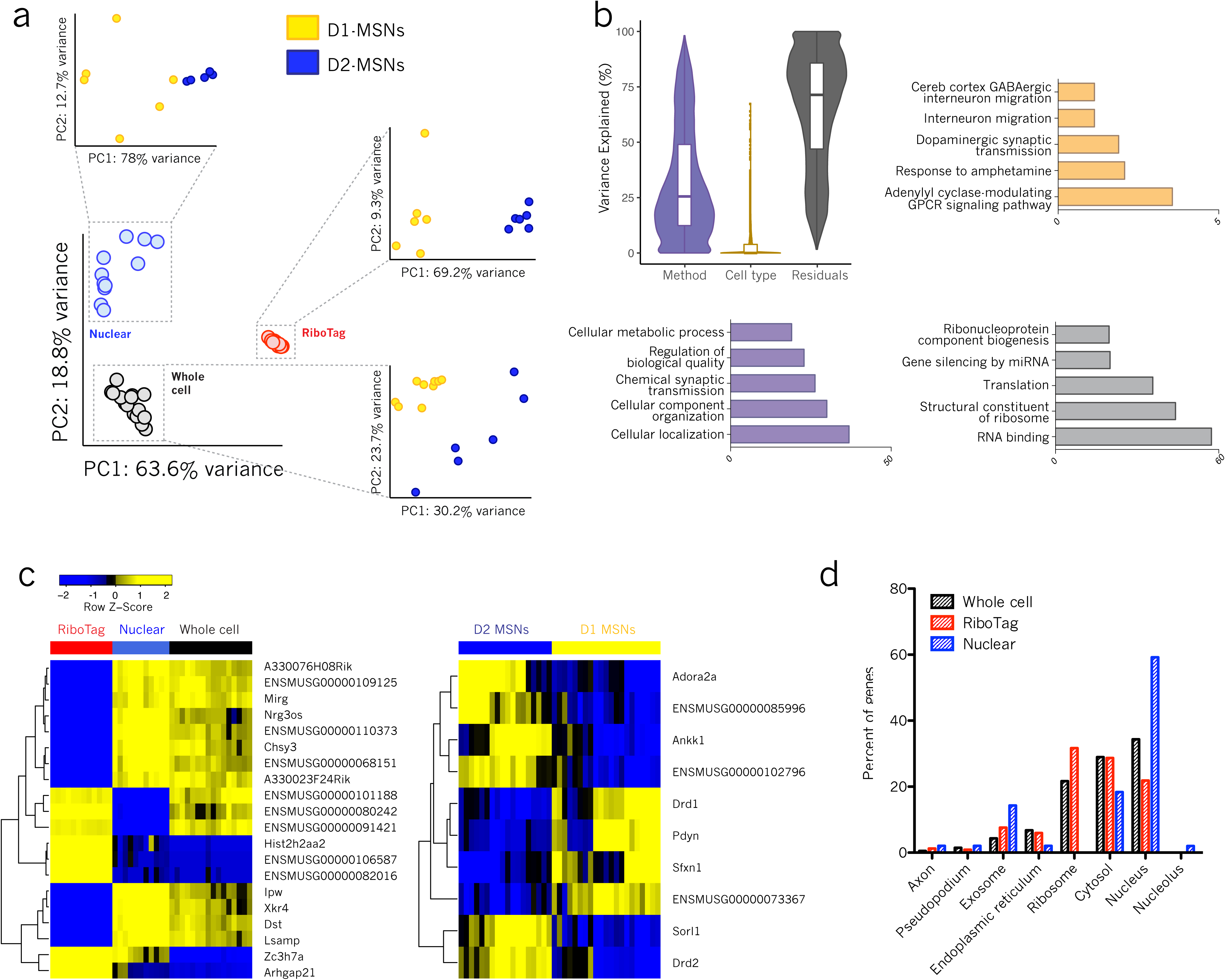
Method accounts for the most variance and method-variable genes are associated with their expected subcellular compartments. (A) Principal component analysis (PCA) across all samples showing separation by method; PCA within method showing separation by cell type (insets) (B) Percent variance explained by method, cell type and residuals with GO enrichment of method-, cell type-, and residual-variable genes (C) Gene expression profiles of the top 20 and top 10 most method- and cell type-variable genes, respectively (D) Subcellular localization of method-variable genes shows enrichment of nuclear-variable genes in the nucleus, and RiboTag-variable genes in the ribosome, as expected

Variation within the total data was further explored using variance partitioning analysis (Fig. 1b). Variance in expression of a given gene that was not explained by method of separation or by cell type was categorized as ‘residual’. On average, 65.3% of transcriptome-wide variance is explained by residual factors, which could include parameters such as the duration and method of trituration^36–38^. This number goes down to 53.6% when only analyzing genes found in all datasets (Supplementary Fig. 3c, Supplementary Table 5). Method accounted for 32.0% of transcriptional variance, and such method-variable genes enriched for expected GO terms relating to cellular localization, which likely has to do with the range of biotypes represented in this gene list (Supplementary Table 6). Our biological variable of interest (cell type) exerted a small effect on transcriptional variance across samples, with only 2.7% of variance explained by cell type (Fig. 2b). This is consistent with the fact that D1- and D2-MSNs are highly similar neurons as noted earlier, which in fact cannot be readily distinguished by PCA on single cell RNAseq of whole striatum^39^. Interestingly, Gene Ontology (GO) analysis of this small list of cell type-variable genes using g:Profiler^40,41^ showed enrichment for expected functions relating to dopamine signaling (Fig. 2b, Supplementary Table 7). This striking – though unsurprising – result highlights the importance of normalizing the method of cellular separation in order to reduce noise in a given dataset and magnify the focus on biological variability.

The top 20 method-variable genes are displayed along with the z-score of their library-normalized expression across method (Fig. 2c). It is interesting to note that many of these top method-variable genes are uncharacterized (8/20, 40%). The percentage of annotated genes steadily increased as the list length increased, demonstrating a bias towards unannotated genes among the most highly method-variable transcripts.

Whole cell- and nuclear-variable genes largely cluster together (Fig. 2c). RiboTag-variable genes comprise a largely non-overlapping pool, though some are shared with whole cell. Future experiments using only one of the methods included here, but seeking to discover consistent biology across cellular compartments, can reduce the technological bias in their data by removing method-enriched genes identified in this study (Supplementary Table 8).

Using RNAlocate, a database of subcellular RNA transcript localization generated across tissues and biological timepoints^42^, we investigated cellular distribution of method-variable genes (Fig. 2d). Our results show the expected subcellular distribution of these genes based on the compartments for which each method respectively enriches. We demonstrate the bias of nuclear isolation towards nuclear-located transcripts, and RiboTag purification towards ribosomal transcripts, with whole cell showing an intermediate profile. Interestingly, the nuclear-variable list showed the highest percentage of exosomally-located transcripts.

### Method-associated topological networks

Variance partitioning analysis revealed genes whose transcriptional variability was determined most by method and cell type. Because many of the method-variable genes were unannotated, potential for further biological interpretation of them was limited. Therefore, in order to generate more functional conclusions about method-related genes, we employed Weighted Gene Co-expression Network Analysis (WGCNA)^43^, which generates transcriptional networks based on the co-expression of genes across a population of samples. For every network module calculated with WGCNA, the eigengene (i.e., first principal component) was used to capture the relative expression changes observed in each module across samples. These eigengenes were further clustered and correlated with each other as well as cell type, method, and/or sex.

Network modules were generated by collapsing across all methods, which resulted in 9 unique modules. Of these 9, correlation analysis discovered one RiboTag-associated module, one nuclear-associated module, and one whole cell-associated module. The top 20 hub genes and their first level connections were extracted in order to generate “hub networks” for modules of interest (see Methods), and these hub networks were probed for functional and biological enrichments using g;Profiler (Fig. 4).

**Figure 3.**
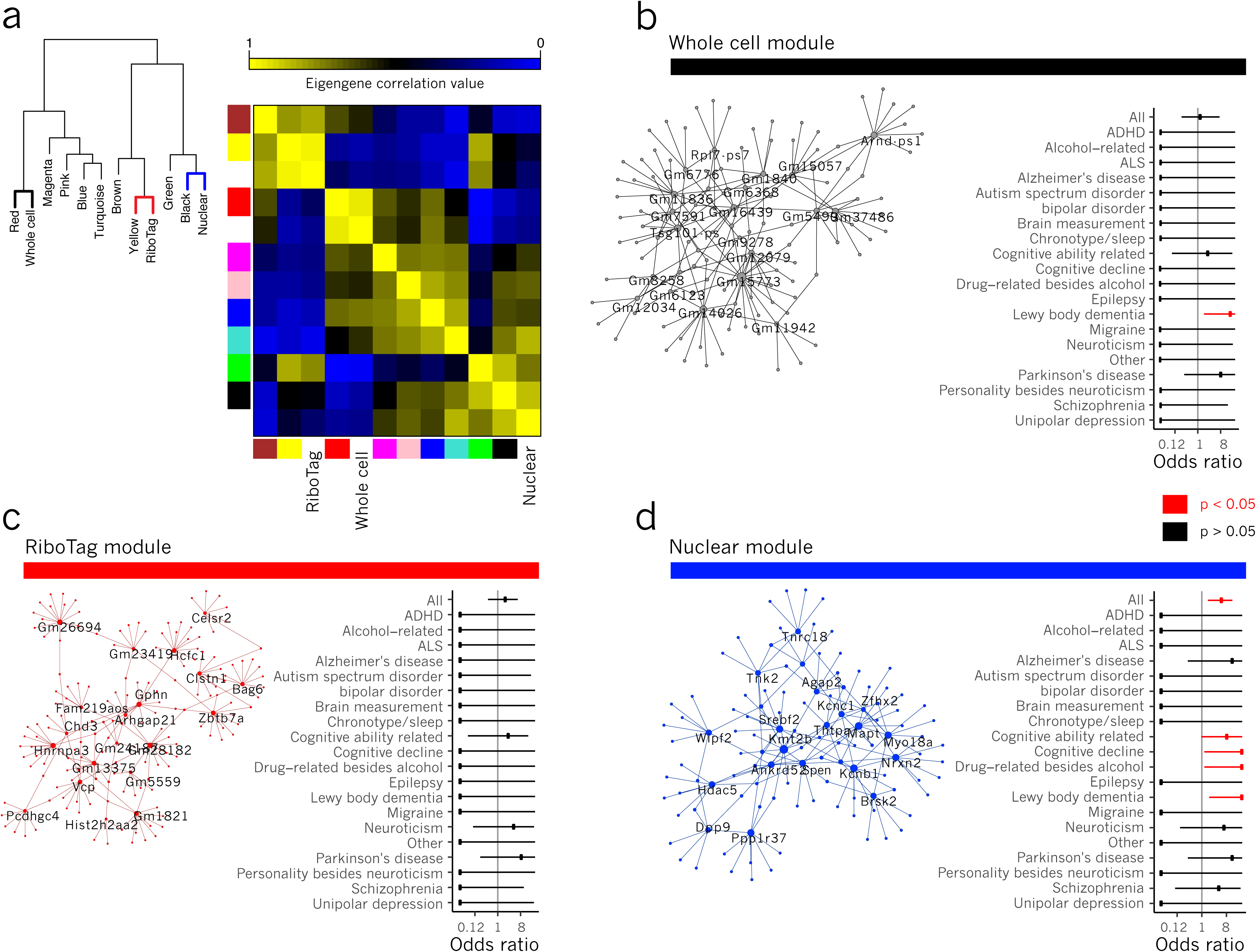
Method-correlated WGCNA modules provide insight into biological function of key regulatory and unannotated genes. (A) Dendrogram showing hierarchical relationship and cluster-map showing correlation values of module and method eigengenes (B) Whole cell hub network including top 20 hub genes (labeled) and their first degree edges; enrichment of GWAS genes among whole cell hub network genes (Fisher’s Exact Test) (C) RiboTag hub network including top 20 hub genes (labeled) and their first degree edges; enrichment of GWAS genes among ribosomal hub network genes (D) Nuclear hub network including top 20 hub genes (labeled) and their first degree edges; enrichment of GWAS genes among nuclear hub network genes

**Figure 4.**
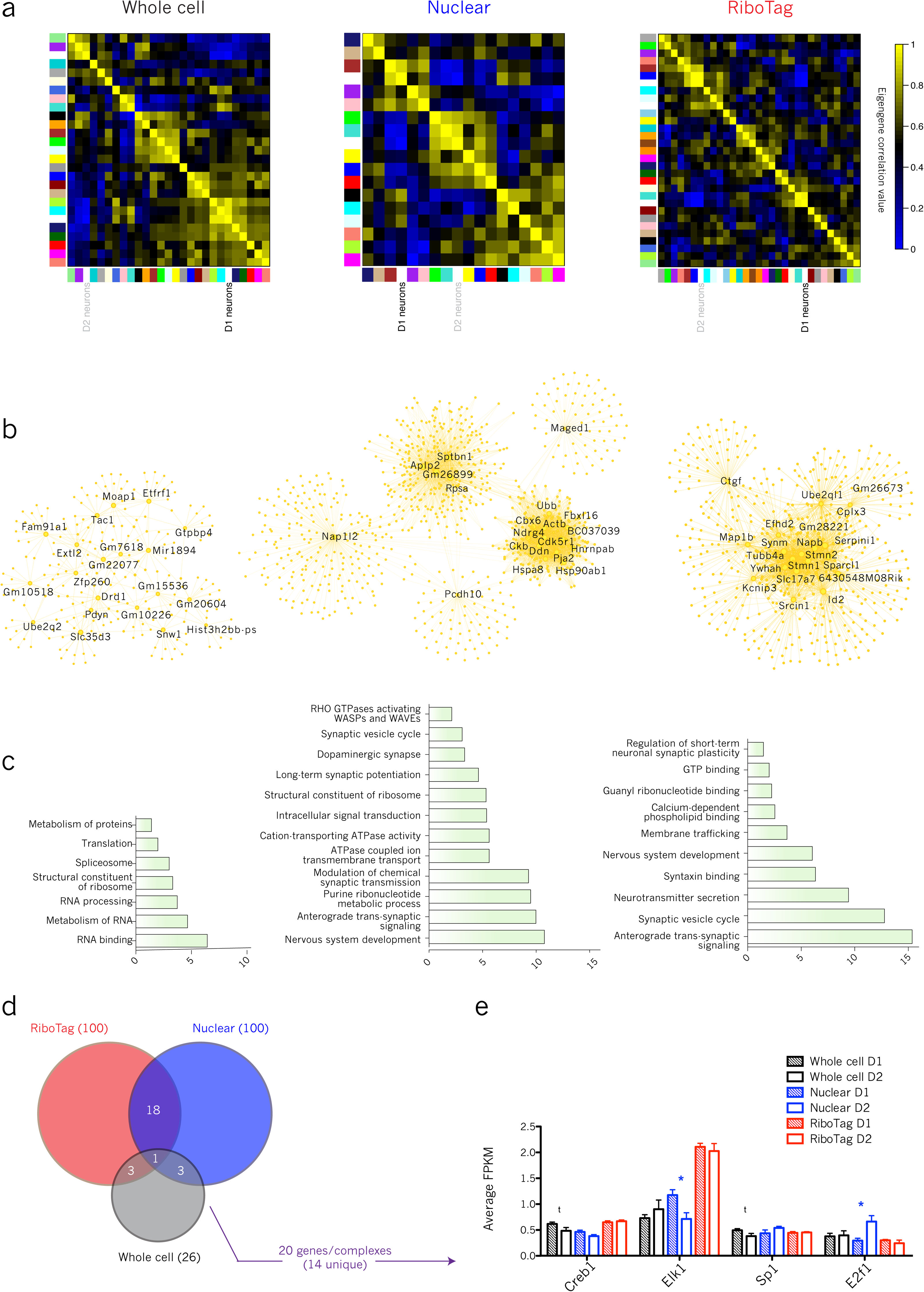
D1 hub networks from multiple methods demonstrate convergent biological function and transcriptional regulation. (A) WGCNA cluster-map with correlations among module eigengenes and D1 cell type; D1 was most correlated with the cyan module in whole cell (Pearson correlation coefficient = 0.91; p-value = 7.0 x 10^-7^), the brown module in nuclear (corr = 0.81; p-value = 2.0 x 10^-3^), and the turquoise module in RiboTag (corr = 0.94; p-value = 8.0 x 10^-6^) (B) D1 module hub networks for each method with hub nodes labeled (C) D1 module GO and KEGG/reactome pathway terms with –log(p-value) of enrichment (D) Overlap of top 100 whole cell, nuclear and RiboTag TFBSs with number of corresponding transcription factors indicated above the arrow (E) Expression (average RPKM) of D1-unique transcription factors across method (asterisk indicates differential expression, pAdj ≤ 0.05)

The RiboTag-associated module enriched for terms relevant to protein-coding activity (GO:0005515, p = 1.48 x 10^-14^; GO:0006412, p = 9.02 x 10^-4^; KEGG:03010, p = 2.21 x 10^-3^; REAC:R-MMU-72702, p = 1.29 x 10^-2^) (Supplementary Table 7). Many of the top twenty hub genes in this network are involved in crucial neuronal functions: organization of DNA (*Hist2h2aa2, Zbtb7a, Hcfc1, Hnrnpa3, Chd3*), regulation of cell growth (*Celsr2, Clstn1*), microtubular organization (*Pcdhgc4, Gphn*), and protein quality control (*Vcp, Bag6*). Their centrality in the RiboTag-associated hub network is congruent with their high levels of expression (all above the 70^th^ percentile of expression by FPKM across method) and their importance in baseline cellular functioning.

The whole cell module enriched for terms relating to cellular energetics (GO:0022900, p = 3.52 x 10^-6^; GO:0006119, p = 1.14 x 10^-5^; GO:0044265, p = 3.61 x 10^-3^) and to RNA processing (GO:0003723, p = 6.04 x 10^-5^; REAC:R-MMU-72163, p = 1.12 x 10^-2^). In comparison, the nuclear module enriched for behaviorally relevant terms (GO:0007610, p = 8.17 x 10^-4^; GO:0050890, p = 4.57 x 10^-3^) and “Voltage-gated potassium channels” (REAC:R-MMU-1296072, p = 1.26 x 10^-2^), highlighting the importance of transcriptional control as a means of regulating K^+^ channel activity^44–46^, as well as perhaps the function of nuclear K^+^ channels in transducing signals of neuronal activity to the nucleus^47,48^.

Using data from the NHGRI-EBI GWAS Catalog^49^, we overlapped hub networks with known risk loci for neuropsychiatric diseases (Supplementary Table 9). In the nuclear hub network, we found a significant overall enrichment (OR = 5.36, Fisher’s exact test two-sided p = 3.23 x 10^-3^, 5/146 term genes), as well as specific enrichments for Lewy body dementia (OR = 145.98, p = 0.014, 1/2 term genes), non-alcoholic drug-related phenotypes (OR = 73.38, p = 0.020, 1/3 term genes), cognitive decline (OR = 73.38, p = 0.020, 1/3 term genes), and cognitive ability (OR = 8.46, p = 0.026, 2/37 term genes). Neither of the other two method-associated hub networks showed overall enrichment for neuropsychiatric GWAS hits, and the only other specific enrichment was in whole cell for Lewy body dementia (OR = 137.89, p = 0.014, 1/2 term genes). This demonstrates an ability of the nuclear method, over and above the other two, to detect disease-relevant genes.

### Cell type-associated topological networks

To investigate the ability of each method to characterize these biologically similar cell types, we performed WGCNA separately on samples from each method. Within each method, hub networks were derived, as above, for modules most correlated to cell type eigengenes. These networks are displayed in Fig. 4 and 5, with node size scaled for relative, within-network degree of connectivity, and hub genes labeled.

**Figure 5.**
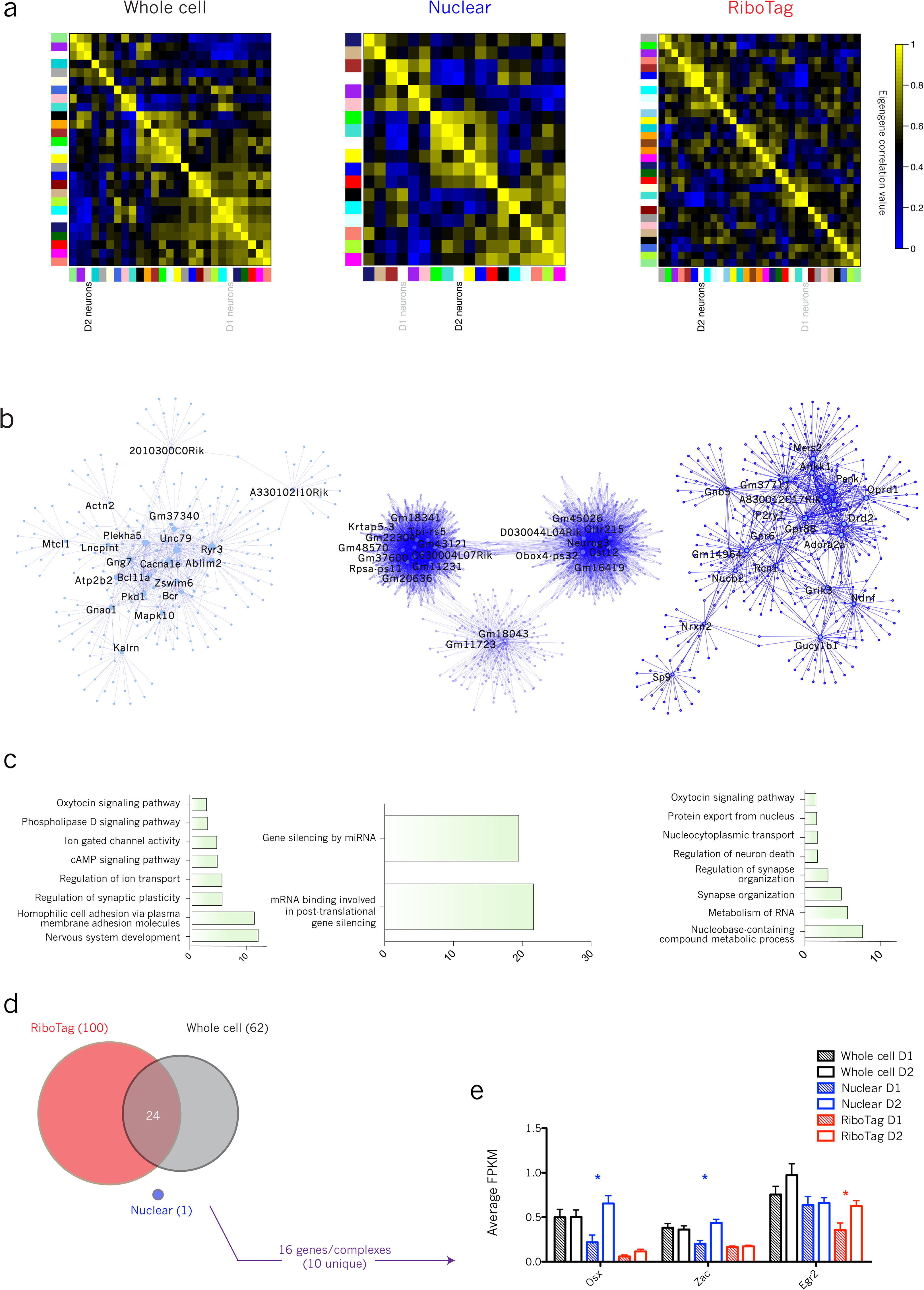
D2 hub networks from multiple methods demonstrate convergent biological function and transcriptional regulation. (A) WGCNA cluster-map with correlations among module eigengenes and D2 cell type; D2 was most correlated with the purple module in whole cell (corr =0.83; p-value = 7.0 x 10^-5^), the turquoise module in nuclear (corr = 0.86; p-value = 6.0 x 10^-4^), and the blue module in RiboTag (corr = 0.98; p-value = 6.0 x 10^-8^) (B) D2 module hub networks for each method with hub nodes labeled (C) D2 module GO and KEGG/reactome pathway terms with –log(p-value) of enrichment (D) Overlap of top 100 whole cell, nuclear and RiboTag TFBSs with number of corresponding transcription factors indicated above the arrow (E) Expression (average RPKM) of D2-unique transcription factors across method (asterisk indicates differential expression, pAdj ≤ 0.05)

Despite a small overlap in hub network genes across methods for D1-MSNs (73/1622, 4.5%) and D2-MSNs (22/1713, 1.3%) (Supplementary Fig. fa), convergent conclusions can be made from the separate techniques. These hub networks were characterized using g:Profiler (Supplementary Table 7) and significantly enriched GO, KEGG and Reactome pathway terms are displayed (Fig. 4c, Fig. 5c). Overlap of genes predicting the same term in separate hub networks is much higher than overall overlap of network genes. For example, both nuclear and RiboTag D1 modules enrich for “anterograde trans-synaptic signaling”, and 12.2% (12/98) of the predictive genes overlap across the two methods.

The top 100 transcription factor binding sites (TFBSs) by enrichment p-value were selected and overlapped for each method. 25 of the top TFBSs for D1 hub networks overlapped, and these corresponded to 20 transcription factors, 14 of which were unique to D1 modules (Fig. 4d). Elk-1 is enriched in D1 nuclei (two-sided t-test, p = 0.02, Fig. 4e), and Creb1 and Sp1 show a trend towards enrichment in D1 whole cells (t-test, p = 0.09, 0.08, Fig. 4e). The fact that not all predicted transcription factors are enriched in the expected cell type – in fact, E2f1 is surprisingly enriched in D2 nuclei despite our upstream analysis indicating it as D1-unique (two-sided t-test, p = 0.01) – can likely be explained either by the inability of upstream analyses to differentiate between multiple members of a transcription factor family or by genome architectural differences between D1- and D2-MSNs, which must be mapped out in order to fully understand the transcriptional dynamics of these cell types.

D1 hub networks generated several terms relating to synaptic function (nuclear, GO:0098916, p = 1.03 x 10^-10^, GO:0060291, p = 2.39 x 10^-5^; RiboTag, GO:0098916, p = 4.36 x 10^-16^; GO:0099504, p = 1.67 x 10^-13^; GO:0007269, p = 3.83 x 10^-10^) (Fig. 4c, Supplementary Table 7). The combined data implicate a D1-enriched function in regulation of synaptic dynamics involving second messenger signaling from the synapse and regulation of vesicle formation and release.

24 of the top 20 TFBSs for D2 whole cell and RiboTag hub networks overlapped, corresponding to 6 overall and 2 D2-unique transcription factors, though none overlapped with the one site predicted from the nuclear dataset (Fig. 5d). However, 6 of the 30 predicted TFBSs for nuclear D2-DEGs did overlap with whole cell and RiboTag module TFBSs (Supplementary Fig. 4b). This suggests that upstream analysis of nuclear sequencing datasets with low average expression levels (Fig. 1e) should be performed on differential expression rather than on network analyses, given the high level of gene co-expression that the nuclear method uncovers. In fact, upstream TFBS analysis on nuclear D1-DEGs also improves overlap of predicted sites (Supplementary Fig. 4c). Despite the failure of the nuclear method to predict overlapping TFBSs, it did most reliably show enrichment by FPKM of these factors in D2-MSNs. Osx and Zac showed significant enrichment in D2-MSN nuclei (two-sided t-test p = 0.006, 0.001, Fig. 5e). RiboTag FPKMs showed enrichement of Egr2 in D2 cells (two-sided t-test, p = 0.02).

Nuclear and RiboTag D2 hub networks showed enrichment for functional terms relating to RNA processing (Fig. 5c, Supplementary Table 7), implicating post-transcriptional mechanisms in the maintenance of D2-MSN identity (nuclear, GO:0035195, p = 5.72 x 10^-25^, GO:1903231, p = 4.27 x 10^-27^; RiboTag, GO:0003723, p = 6.16 x 10^-13^; REAC:R-MMU-8953854, p = 2.12 x 10^-6^). The importance of post-transcriptional regulation in D2-MSNs helps to explain the higher variance observed in D2 compared to D1 MSN nuclei (Supplementary Fig. 5).

### Analyzing sex differences using RiboTag-RNAseq

The RiboTag-RNAseq dataset was considerably less complex than those generated by the other two methods, and therefore showed the most robust separation of cell types on PCA. We thus expected it to most powerfully separate two biological types – namely, MSNs from male and female mice – whose difference has not yet been characterized. We compared the aforementioned male mice to 10 RiboTag-RNAseq samples (D1 n=5, D2 n=5) from female mice. WGCNA performed on RiboTag-purified male and female D1**-** and D2-MSN RNA generated modules (Fig. 6b, 6c), which correlated with the eigengenes for male and female sex. Notably, the correlation p-value for the male-enriched module did not meet statistical significance (Pearson correlation p = 0.2; female, p = 0.008). This suggests that MSN differences by sex are less pronounced than differences between cell types within a given sex. Nevertheless, male-and female-correlated modules were both enriched for terms relating to metabolism of genetic material (p < 9.6 x 10^-2^), a result mirrored in GO analysis of previous whole tissue RNAseq^50^ (Supplementary Table 7). The female hub network predicted Tel1, a regulator of telomere maintenance, as an upstream transcription factor (Fig. 6c). This prediction is in line with evidence of sexual dimorphism in telomeric structure^51,52^, and demonstrates that sex differences in the brain can be investigated at the cell type-specific level. This is possible even when cell type-enriched DEGs are highly overlapping across sex (Fig. 6d).

**Figure 6.**
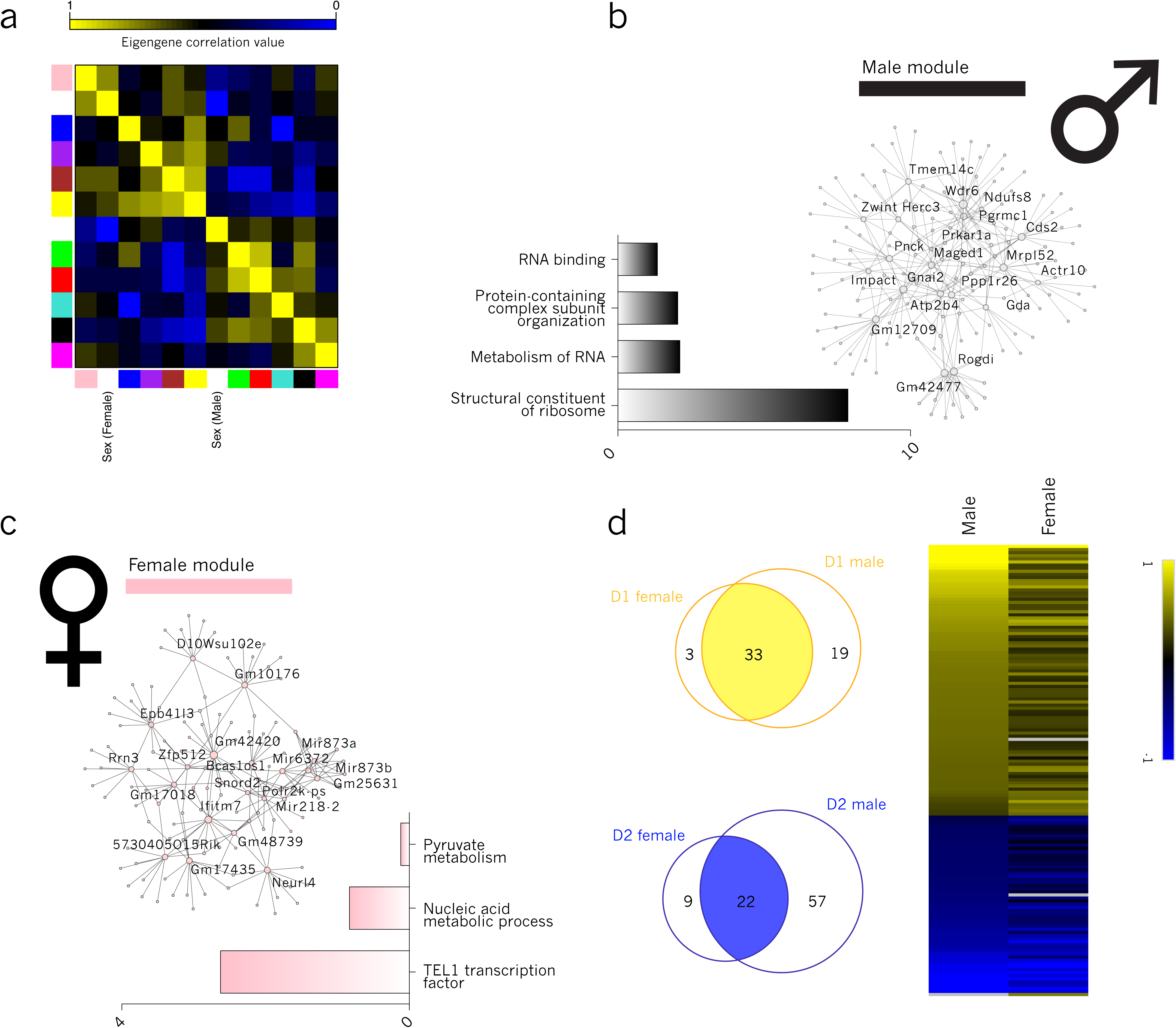
RiboTag data can discern small-variance biological variables, with WGCNA generating sex-correlated modules.

(A) RiboTag WGCNA cluster-map showing correlation values of module eigengenes and sex; male sex was most correlated with the black module (corr = 0.27; p-value = 0.2) and female sex was most correlated with the pink module (corr = 0.55; p-value = 8.0 x 10^-3^)

(B) Black module hub network (male) with hub nodes labeled; GO and pathway terms with –log(p-value) of enrichment displayed

(C) Pink module hub network (female) with hub nodes labeled; GO terms and TFBS prediction with –log(p-value) of enrichment displayed

(D) Overlap of D1- and D2-DEGs from male and female datasets; union heatmap of these DEGs showing log2foldchange (D1 vs. D2) in each dataset (black = low fold change, grey = not detected in the dataset)

## Discussion

RNAseq is a powerful method for examining the baseline, homeostatic mechanisms that determine cellular identity, but its output is highly dependent on the method by which libraries are prepared. We examined, in particular, the effect of cell separation technique on RNAseq of biologically similar populations of dopamine receptor-expressing D1- and D2-MSNs in the NAc. Our results highlight, foremost, the importance of reducing technique-induced noise by implementing well-controlled experiments and using strict FPKM minimum and maximum cutoffs to eliminate sequencing artifacts.

This work also provides several gene lists that will inform future work. Our modeling of transcriptional control mechanisms can be used to make further predictions about the subcellular transcriptional regulation of various transcripts. Our variance partitioning data present a resource of technique-variable genes with which to normalize when comparing datasets generated using different cell separation techniques. As well, our findings expand on existing RNA localization datasets by demonstrating a robust list of genes expressed only in the nuclear and whole cell datasets, including genes whose transcripts are restricted to the nuclear compartment.

We confirm the bias of RiboTag affinity purification towards isolation of protein-coding genes, a fact which affords both advantages and disadvantages. The disadvantage is, of course, the failure of this technique to recover large sets of non-coding regulatory genes which differ dramatically between cell types (as indicated by nuclear and whole cell sequencing) and are known to play an important role in biological regulation. Its advantage is the superior separation of cell types as compared to the other methods. The RiboTag dataset showed the clearest separation of D1- and D2-MSNs, although it also showed the lowest fold-change in differential expression of cell type-enriched genes.

Nuclear- and whole cell-FACS-based isolation similarly have advantages and disadvantages. One challenge is their extreme complexity – strict cutoffs must be set, especially when using the nuclear method, in order to focus on only the most robust results. This complexity is furthered by the presence of immature mRNAs, which may or may not be destined for translation. For cell types whose nucleus contains lowly expressed, high variance genes, nuclear sequencing generates noisier results than either whole cell or RiboTag sequencing, and upstream analysis of differential expression, as opposed to co-expression data, generates more reliable conclusions.

However, the nuclear method does offer advantages. A methodological advantage of nuclear isolation is the fact that it can be performed on frozen tissue, making implementation of large experiments and/or use of banked human tissues much more manageable. A conceptual advantage of the nuclear method is demonstrated by our GWAS hit enrichment analysis, which reveals that this method captures more disease-relevant loci than either of the other two, making it an appealing option for those studying the genetic basis of neuropsychiatric disease.

Despite these differences between methods, and the relatively low overlap of differentially expressed genes, convergent conclusions can be made about cell type-enriched transcription. We were able to robustly identify regulatory transcription factors, central biological functions, and enriched protein complexes across methods.

An important future question concerns the ability of these techniques to capture transcriptional response of cell populations to various stimuli. We would expect nuclear and RiboTag-generated datasets to represent different stages of response, and to be differentially valuable during given post-stimulus time windows. This is a question which requires empirical investigation, and to which baseline normalization based on our results can be applied.

## Methods

### Transgenic animal lines

All animal protocols were approved by IUCAC. All mice were bred on a C57BL/6J background. Mice used for nuclear isolation heterozygously expressed a nuclear GFP label under the promoter of either Drd1 or Drd2 (Drd1-cre x Thy1-^loxP^STOP^loxP^-EGFP-F; Drd2-cre x Thy1-^loxP^STOP^loxP^-EGFP-F). Mice used for whole cell isolation heterozygously expressed a cytoplasmic fluorophore under the promoter of either Drd1 or Drd2. Mice used for RiboTag affinity purification expressed an HA-tagged ribosomal subunit under the control of Drd1 and Drd2 (Drd1-cre x Rpl22-^loxP^exon4^loxP^-exon4-HA; Drd2-cre x Rpl22-^loxP^exon4^loxP^-exon4-HA).

### Cell isolation

Each sample represents tissue from a unique mouse. Nuclear samples were obtained from frozen tissue. This tissue was mechanically dissociated and nuclei lysed using a glass douncer in ice-cold lysis buffer (10.94% w/v sucrose, 5 mM CaCl_2_, 3 mM Mg(CH_3_COO)_2_, 0.1 mM EDTA, 10 mM Tris-HCl pH 8, 1 mM DTT, in H_2_O). Samples were centrifuged and supernatant removed and placed on top of a 60% sucrose solution (60% w/v sucrose, 3 mM Mg(CH_3_COO)_2_, 10 mM Tris-HCl pH 8, 1 mM DTT, in H_2_O). The sucrose gradient was centrifuged at 24,400 rcf for an hour, and nuclei at the bottom of the gradient were resuspended in PBS. DAPI was added at a concentration of 0.5 μg/mL. Whole cell samples were obtained from fresh tissue, which was rotated at 37°C for 45 minutes in 1 mg/mL papain suspended in digestion buffer (5% w/v D-trehalose, 0.05 mM APV, 0.0125 mg/mL DNAse, in Hibernate^TM^-A (Thermofisher, A1247501)). Tissue was then placed in FACS buffer (0.58 mg/mL albumin inhibitor (Worthington Biochemical, LK003182), 5% w/v D-trehalose, 0.05 mM APV, 0.0125 mg/mL DNAse, in Hibernate^TM^-A) and triturated using progressively smaller pipette tips. Samples were passed through a 70 μm filter and placed on top of a layer of 10 mg/mL ovomucoid-albumin in FACS buffer. The pellet was resuspended in FACS buffer and DAPI was added at a concentration of 0.5 μg/mL. RiboTag samples were obtained from fresh tissue as previously described^53,54^.

### Library preparation and sequencing

All isolated RNA was prepared using the Clontech SMARTer^®^ Stranded library preparation kit (cat. no. 634838). Briefly, samples were ribo-depleted, and total RNA was reversed transcribed, classified using Illumina indices and amplified with 8 cycles of PCR. Samples were sequenced to a minimum depth of 30 million reads using paired-end reads with V4 chemistry on an Illumina Hi-Seq machine.

### RNAseq alignment preprocessing

Reads were aligned to GRCm38 with HISAT2^55^. All aligned samples were reviewed for quality control with FASTQC (see URLs). Reads were counted for Mus_musculus.GRCm38.90 using featurecounts with the settings strandSpecific=0 allowMultiOverlap=T, countMultiMappingReads=F, and isPairedEnd=T^56^. Gene level counts were generated from both only exons and all features. The percent of introns per gene was obtained by subtracting gene-level exon counts from all counts. Gene-level counts from all features were used unless otherwise specified.

Data were processed in three tranches: (1) male samples from all methods pooled, (2) male samples within each method, and (3) male and female RiboTag samples pooled. Genes with >1 fragments per million in ≥2 samples were kept for initial analyses. Final filters were ≥1 fragments per million per kilobase (FPKM) in at least one sample for nuclear and whole cell, and ≥0.1 FPKM in at least two samples for RiboTag. Counts were normalized to effective library size, and variance stabilized with DESeq2 b Design matrix covariates, depending on the tranche, method (whole cell, RiboTag, or nuclear), cell type (D1 or D2) and/or sex (male or female). Variance stabilized transcripts (from the DESeq2 vst function) were used for both principal component analyses (PCA) and weighted gene co-expression network analysis (WGCNA)^57^. Two RiboTag samples were excluded because they were PCA outliers and had >80% of reads unmapped.

### Differential expression

Differential expression was assessed between D1 and D2 MSNs within each method using DESeq2^58^. To determine cell type-enriched genes, lists were filtered for |log_2_foldchange| > 0.38 corresponding to a 30% change (past work has demonstrated that at least a 15% change is needed to replicate with qPCR^59^), pAdj ≤ 0.05, mean normalized counts above the 25^th^ percentile within method. Nuclear data were also filtered for log_2_foldchange standard error below the 75^th^ percentile. Length and biotype data for each gene were obtained using the R package biomaRt^60^.

### Variance partitioning

Euclidean distance was calculated from PC1 and PC2. Centroids were determined from the mean PC values within each method. Distance was calculated with Euclidean distance from the cluster centroid, and pairwise t-tests between all groups were performed for Euclidean distances.

To calculate the percent variance explained by every covariate for every gene, VariancePartition was run on all genes and the intersection of genes passing minimum detection thresholds within each method^61^.

Based on the percent variance explained, genes were assigned to one of three covariate categories (method, cell type, residual). Genes within each category were analyzed using g:Profiler, and five of the top ten most significantly enriched terms are displayed. Method-variable genes were further divided by enrichment within method. Genes were assigned to the method in which they had the highest RPKM, and overlapped with the RNAlocate database. Genes in our analysis that were not represented in the RNAlocate database were removed from the subcellular localization analysis.

### High confidence co-expression networks

Networks were generated in three tranches specified previously, all male samples, separately for each method, and for RiboTag with both male and female. DESeq2 variance stabilized transcript expression matrices were used as input.

For WGCNA, soft power thresholds for signed correlations were chosen to achieve approximate scale-free topology (R^2^ > 0.8): β_AllMale_=18, β_Nuclear_=12, β_WholeCell_=18, β_RiboTAG_=9, β_RiboTAGwithFemale_=6. The WGCNA function blockwiseModules was run with the appropriate power and the parameters networkType = “signed”, TOMType = “signed”, detectCutHeight = 0.99, minModuleSize = 100, reassignThreshold = 0, minKMEtoStay = 0.1, mergeCutHeight = 0.2, corType=“bicor”, numericLabels = TRUE, and pamStage = FALSE. Networks were then plotted and modules were exported as edge and node files from the topological overlap matrix.

ARACNE networks were generated using the minet R package by first building a mutual information matrix (build.mim function with estimator = “mi.empirical” and disc = “equalfreq”)^62,63^.

Edges detected independently with both WGCNA and ARACNE were kept, and these edges were used to determine the top 20 hubs per network (by degree) as well as their first degree neighbors.

Genes within each network were analyzed using g:Profiler, and representative terms from those most significantly enriched are displayed.

### GWAS gene set enrichment

The GWAS catalog v1.0.2 (release 2018-08-14) was downloaded from the NHGRI-EBI catalog (see URLs)^49^. Associations were kept if the lead variants had p<5×10^−8^ and mapped within a gene’s start and stop coordinates (*i.e.*, was not intergenic). The gene associated with the lead variant was then mapped to its ENSEMBL mouse orthologue (see URLs)^64^. The remaining 1843 traits were manually curated for neuropsychiatric relevance, and categorized into 20 superseding traits, resulting in a final set of 554 genes from 733 gene-trait pairs.

Gene set enrichment between GWAS genes and network hubs was calculated from a 2×2 contingency table with a Fisher’s Exact Test. The contingency table comprised all genes used as input for WGCNA and ARACNE for a given tranche, with rows labeling a gene as GWAS or not and columns labeling genes as belonging to a given hub network.

All custom code is available under the MIT license (see URLs).

### URLS

Custom code: https://github.com/frichter/d1_d2_rnaseq/

FASTQC: https://www.bioinformatics.babraham.ac.uk/projects/fastqc/

GWAS catalog: https://www.ebi.ac.uk/gwas/docs/file-downloads

Human-mouse orthologues: http://useast.ensembl.org/biomart/martview/

## Author information

### Acknowledgements

This work was supported in part through the computational resources and staff expertise provided by Scientific Computing at the Icahn School of Medicine at Mount Sinai. Also at the Icahn School of Medicine at Mount Sinai, this work was supported by the facilities of and guidance from the Flow Cytometry core.

### Author contributions

F.R. and H.K. analyzed and interpreted results and wrote the paper. H.K., B.L., and R.C. generated the data. All authors discussed the analysis and results and commented on the manuscript

### Competing interests

The authors have no competing interests.

## Supplementary figure legends

**Supplementary Figure 1. Differences in genomic distribution and differential expression across methods.**

(A) Percent of intronic reads across samples for each gene, median indicated as a line in method color
(B) Differential expression overlap using lenient filters on all (1 FPM in at least two samples)
(C) Differential expression overlap using strict filters on all (FPKM ≥ 1 in at least one sample and all samples with FPKM < 5.0 x 10^4^)
(D) Differential expression overlap using strict filter for whole cell and nuclear and a less strict filter for RiboTag (FPKM ≥ 0.1 in at least two samples and all samples with FPKM < 5.0 x 10^4^)
(E) Density plots of gene lengths for all genes, protein-coding genes, and non-coding genes for each method (whole cell = black, nuclear = blue, RiboTag = red)

**Supplementary Figure 2. Hub network overlaps and nuclear DET-predicted TFBSs**

(A) Overlap of hub network genes for D1- and D2-correlated modules across method
(B) Overlap of genes showing regulation by transcription, cytosolic mechanisms, or translation in D1- and D2-MSNs. 0 overlaps are not labeled.
(C) List of 27 miRNAs predicted upstream of the (Transcription D1) / (Cytosol D2) overlap (overlap p-value < 0.05) and FPKM values of the 4 miRNAs from this list that are differentially expressed in D2-nuclei (but not in D2-whole cells)

**Supplementary Figure 3. Variance in just genes detected in all three methods.**

(A) PCA of combined data showing separation of samples by cell type on PC5/PC6. Key is shown in figure
(B) PCA of nuclear, whole cell, and RiboTag datasets from this study along with RiboTag RNA-sequencing from Song et al, 2018. Key is shown in figure
(C) Variance explained by method, cell type and residuals across the transcriptome. Central bars represent median

**Supplementary Figure 4. Hub network overlaps and nuclear DET-predicted TFBSs**

(D) Overlap of hub network genes for D1- and D2-correlated modules across method
(E) Overlap of predicted TFBSs for D2 whole cell and RiboTag hub networks with predicted TFBSs for nuclear D2-DEGs
(F) Overlap of predicted TFBSs for D1 whole cell and RiboTag hub networks with predicted TFBSs for nuclear D1-DEGs

**Supplementary Figure 5. Variance distributions for D1- and D2-DETs across methods.**

(A) Variance distribution for nuclear D1- and D2-DEGs (top of the box = 3^rd^ quartile, midline = median, bottom of the box = 1^st^ quartile)
(B) Variance distribution for RiboTag D1- and D2-DEGs (top of the box = 3^rd^ quartile, midline = median, bottom of the box = 1^st^ quartile)
(C) Variance distribution for whole cell D1- and D2-DEGs (top of the box = 3^rd^ quartile, midline = median, bottom of the box = 1^st^ quartile)

## Supplementary table list

Supplementary Table 1: Nuclear DEG logistic regression

Supplementary Table 2: Differential expression by method

Supplementary Table 3: FPKM by method

Supplementary Table 4: FPKM and variance correlations across methods

